# Rats synchronize predictively to metronomes

**DOI:** 10.1101/2023.06.19.545615

**Authors:** Vani G. Rajendran, Yehonadav Tsdaka, Tung Yee Keung, Jan W. H. Schnupp, Israel Nelken

## Abstract

Humans easily and naturally synchronize their motor activity to music, a behavior unparalleled by other species. As a consequence of the mystery surrounding its evolutionary and biological origins, there is rapidly growing interest in exploring the capacity of nonhuman species to synchronize motor activity with auditory rhythms. The hallmark observation in human studies is that synchronization is predictive. Given the highly variable behaviors observed in other animals, it is particularly difficult to distinguish a predictive synchronization behavior from one that perhaps resembles it, but relies on simpler strategies. Here, we introduce a novel modeling approach that quantitatively compares candidate strategies for explaining observed behaviors. Eight rats synchronized to metronomes across a range of tempi (0.5-2 Hz), and immediate water reward was delivered whenever they initiated a lick burst within a short time window around beats. We observed a roughly constant baseline lick probability throughout, with a ∼30% modulation of the lick rate around beats. We fitted to the results six candidate models, ranging from an ‘insensitive’ model assuming that rats lick at random, to predictive models where rats suppress licking between beats for a tempo-dependent duration of time. The predictive model consistently outperformed other models, including a reactive model, demonstrating that rats synchronize predictively to auditory metronomes.

## Introduction

Humans synchronize their movements to auditory stimuli, while nonhuman species largely do not. The origins of musical beat perception and synchronization in humans has remained elusive, prompting a recent surge in studies investigating rhythm perception and synchronization across a variety of nonhuman species. These studies include tests of the ability to discriminate isochronous from non-isochronous rhythms [1-3], metronome tapping [4-8], and studies of spontaneous movements to rhythms and music [9-12]. It has been hypothesized that beat perception is a byproduct of vocal learning ability [13], but this hypothesis needs to be revised because many of the species that show some capacity for rhythm perception and production are not vocal learners. It seems therefore that the auditory-motor coordination required for rhythmic synchronization developed gradually during mammalian evolution [14]. The most commonly used laboratory mammals in neuroscience studies are rodents (rats and mice, non vocal learners). A clear demonstration of predictive synchronization in rodents would vastly expand the range of tools and approaches available towards understanding the neural mechanisms of beat perception and synchronization.

Indeed, recent studies in macaques (a non vocal learner) demonstrate the depth of insight that can be gained through in-vivo recordings during synchronization to simple periodic beat sequences, or metronomes [15, 16]. Interestingly, while macaques do not spontaneously synchronize to auditory rhythms, they are nevertheless able to produce human-like predictive tapping behaviors with appropriate training [6, 17]. Similarly, rats are able to discriminate isochrony from non-isochrony [2] and even recognize a familiar rhythm from an unfamiliar one [18]. Neural responses in rodents to auditory rhythms and music also reflect general sensory processing mechanisms that may predispose beat perception [19-21]. However, in a metronome tapping task, rats showed a behavior that is far less precise than that of humans and macaques, making it difficult to determine whether the behavior is predictive [7]. A recent study also observed spontaneous head movements in rats while listening to real music [12]. In that study, however, the subtle head movements were present only during the first experimental session, and since the animals were not performing a task, it is difficult to interpret these movements as an unambiguous demonstration of auditory-motor synchronization. While the observations from these studies are highly suggestive, they do not unequivocally demonstrate that rats can predictively synchronize to beat, but neither do they clearly demonstrate otherwise.

Proving whether a synchronization behavior is predictive or not is a nontrivial task, particularly when the behavior differs from the 1-to-1 beat-to-tap correspondence typical of humans. In humans, three main features are taken to indicate prediction: negative mean asynchrony (NMA), negative lag-1 autocorrelation, and Weber law-like increase in errors with larger intervals [22, 23]. These key features of human tapping have been the standard against which animal synchronization studies have been compared, with impressive success in macaques [6, 16].

We argue here that by requiring a human-like 1-to-1 beat-to-tap behavior from animals who may have no ecological necessity or ability to “tap to the beat,” we overlook the possibility that they may nonetheless show predictive motor responses in other ways [24]. To cope with the varied forms that behavioral data from animals may take, we propose here a more general theoretical construct, which we call ‘latent predictive synchronization’ to a rhythm. Latent predictive synchronization is a behavior in which 1) motor responses are temporally locked (potentially only probabilistically) to periodic beats, and 2) the periodic changes in the probability of these motor responses depend on the beat rate. Importantly, requiring both of these conditions avoids confounding actions that are stereotypically locked to beats (e.g. startle reflexes or purely reactive responses to beats), or behaviors that depend on the beat rate but are not locked to beats (e.g. changes in the rate of randomly timed movements that depend on tempo), with predictive synchronization. At the same time, the notion of latent predictive synchronization allows a large degree of flexibility in how a predictive behavior might look. In particular, it allows for a 1-to-many beat-to-action correspondence, but only if the timing of changes in the statistical properties of the elicited actions are both locked to, and depend on, the beat rate.

Demonstrating latent predictive synchronization requires shifting from the standard event-centric (e.g. tap-based) view of rhythmic synchronization, to one that is state-based (e.g. high probability of tap, low probability of tap), noting that regular 1-to-1 beat tapping is simply a special case of this state-based formulation. Here, we developed a Markovian approach that enables latent predictive synchronization to be quantitatively distinguished from other behavioral strategies. In the experiments described here, rats were trained to synchronize their licks to metronome beats for reward. Licking has the advantages of being natural and enabling the delivery of immediate feedback through water reward, but is difficult to volitionally control [25]. The model therefore consisted of two basic states – licking and not licking – with parameters that could be modulated by external events (metronome beats and water rewards). Different candidate strategies used by the animal were modeled by constraining these modulations in different ways, ranging from total independence to beat rate-dependence indicative of prediction. These models were then compared to the data and their goodness of fit determined. Based on data from eight rats licking to metronomes with rates randomly chosen between 0.5 and 2 Hz, and comparing six different candidate strategies, we demonstrate explicitly that rats indeed synchronize predictively to auditory metronomes.

## Results

### Rats lock their licking to metronome beats

The task, shown in Fig. 1, was designed to test whether rats could lick from a water spout in synchrony with an auditory metronome. Each trial in a session consisted of 16 isochronous noise bursts with a constant inter-stimulus interval (ISI). The ISIs were selected randomly for each trial from a log-uniform distribution between 500 ms and 2000 ms (Fig. 1A), and trials were separated by 3-4 s of silence. Rats received immediate feedback on the relative timing (asynchrony) of each lick burst with respect to the nearest sound onset through water reward delivered through the spout they were licking (Fig. 1B). A small drop of water was delivered immediately (<3 ms delay) if a lick satisfied two conditions: it was within a time window of ± 7.5% ISI relative to a sound onset, *and* it was separated from any preceding licks by at least 450 ms (Fig. 1C). The tight requirements for reward availability ensured that continuous licking was not rewarded at all, and incentivized predictive strategies.

**Figure 1.**
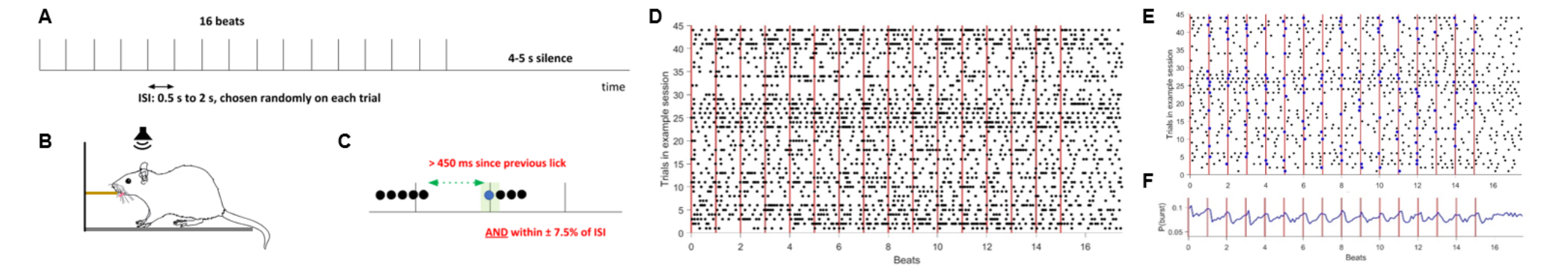
Task design and raw data. **A)** Trials consisted of 16 isochronous metronome beats. The inter-stimulus interval (ISI) for each trial was selected randomly from a log-uniform distribution between 0.5-2s. Trials were separated by 3-4 seconds of silence. **B)** The rat was placed in a custom cage and sound was delivered by a speaker placed above the cage. The rat was trained to lick a water spout in synchrony with metronome beats. **C)** The spout would immediately release a drop of water if a lick satisfied two conditions: that it was within +/-7.5% ISI of a beat onset, and was separated from the previous lick by ≥450 ms. **D)** Raster depicting licks in an example ∼30 min session. Each row is one trial, and since every trial had its own tempo, time is normalized to the ISI of each trial. **E)** Same data as D, but showing only the first lick in each burst. Rewarded lick bursts are marked in blue. **F)** A histogram of lick burst onsets around metronome beats, pooled across all sessions, trials, and animals. Y-axis is the probability of lick burst onset in each 0.1*ISI bin.

Data from a typical session are shown in Figure 1D. Licks were usually produced in bursts with a typical rate of about 7 Hz, consistent with evidence that licking in rats is governed largely by a central pattern generator in the brainstem [25]. With training, however, the rats learned to modulate their licking in order to satisfy the conditions for reward. Figure 1E shows only the onsets (the first lick) of each lick burst, with rewarded lick bursts highlighted in blue. While lick bursts are not produced solely around beat times, a clear modulation of lick burst probability around beats is apparent (Fig. 1F). The results that follow use the onsets of lick bursts as the events of interest, since reward was determined by these first licks when timed correctly.

Figure 2 illustrates the timing of the lick bursts produced by all animals across the range of tempi tested in this task. The time difference between the onset of a lick burst and the nearest sound onset was defined as the lick burst asynchrony. Since every trial had its own tempo, these asynchronies are expressed as phases (centered around beat onsets, range −0.5 to +0.5 of the metronome period).

**Figure 2.**
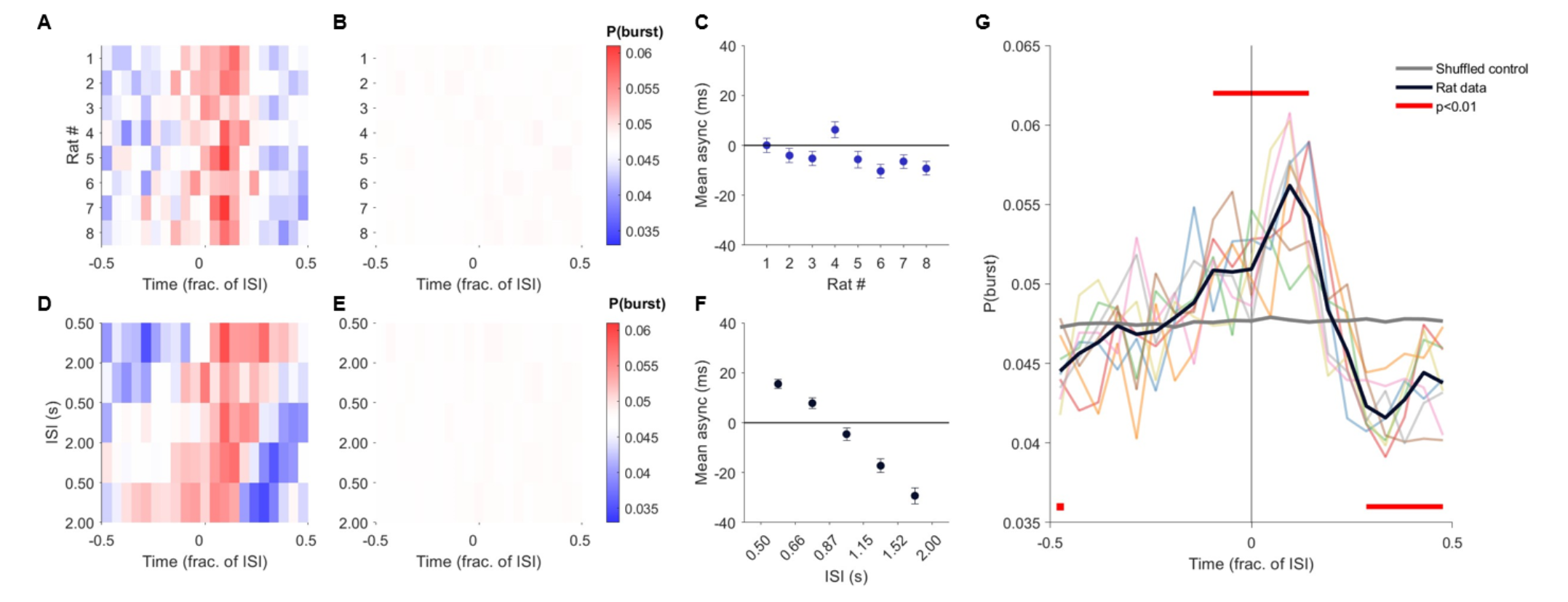
Rats significantly modulate their licking around metronome beats. **A)** Probability of emitting a lick burst over the beat interval (normalized by ISI, −0.5 to +0.5), pooled across all trials of all rates. Individual animals (rows) showed consistent licking patterns. **B)** Shuffled control for A (100 permutations). **C)** Mean asynchronies for individual rats; most are near zero or slightly negative. **D)** Probability of lick burst over the beat interval, pooled across animals and split into five tempo bins. **E)** Shuffled control for D. **F)** Mean asynchronies by tempo bin. Asynchronies become more negative for slower metronome rates. **G)** Grand average probability of lick burst, pooled across all animals and rates (black). Individual animals are shown by the colored lines (same data as in A), the grand average shuffled control is shown in grey. Time bins where lick burst probability is significantly different from the shuffled control are marked in red (p<0.01, permutation test with False Discovery Rate correction using Benjamini-Hochberg step-up procedure, 100 permutations).

The probability of producing a lick burst showed a distinct modulation pattern, increasing before the beat and decreasing following it (Fig. 2A). This modulation was highly consistent across animals. The lick burst asynchronies were significantly modulated by phase for all eight rats (p<10^-5^, circular Rayleigh test for nonuniformity, N=7,000-11,000 lick bursts per rat). To explore whether this pattern could have been observed with random licking, a simple control simulation was performed that preserved the sequence of the observed lick bursts, but analyzed them as though they occurred during another randomly drawn metronome ISI (see *Methods*). The shuffled lick burst asynchronies (Fig. 2B) were not significantly modulated by phase for any of the rats (p>0.6, circular Rayleigh test for nonuniformity, N=6,000-8,000 shuffled lick bursts per rat). The mean asynchronies of lick bursts were on average close to zero or negative for most rats (Fig. 2C).

To explore how lick burst asynchronies varied with tempo, trials were split into five log-spaced tempo bins and pooled across all animals (Fig. 2D). Lick burst probability was significantly modulated by phase in each tempo bin (p<10^-6^, circular Rayleigh test for nonuniformity, a total of N=8,000-24,000 lick bursts from 8 animals per tempo bin). The same analysis performed on the shuffled data abolished the modulation in all tempo bins (Fig. 2E; p>0.3, circular Rayleigh test for nonuniformity, N=8,000- 24,000 shuffled lick bursts per tempo bin). The mean asynchronies (Fig. 2F) reflect the monotonic shift towards more negative values with increasing ISIs that is apparent in Fig. 2D, hinting at a predictive control of lick burst time.

The grand average histogram of the lick rates, pooled across all animals and tempi, is shown in Figure 2G, along with the grand average shuffled control histogram. The overall modulation in lick burst probability observed is approximately 30% of the baseline lick rate. The data show significant deviations from the control at many phases even before the beat, again suggesting predictive modulation of lick probability by the rats.

### Are rats reacting or predicting? A test of six possible strategies

To determine whether the observed patterning of lick bursts reflects latent predictive synchronization, we developed a Markovian framework by which different underlying strategies for shaping lick rates can be modeled and compared. The state graph of the process is shown in Fig. 3A and is shared by all models. The graph has two branches, a ***Burst*** branch that consists of states traversed while the animal is in a lick burst, and a ***Break*** branch that consists of states traversed while the animal is not licking. The states along a branch represent time elapsed while the animal is in a Burst or in a Break. Thus, every time point during a trial is assigned a process state that consists of two elements: the behavioral state of the animal (Burst or Break), and how long the animal has already been in that behavioral state since the last transition (represented by the location along the branch, Fig. 3B).

**Figure 3.**
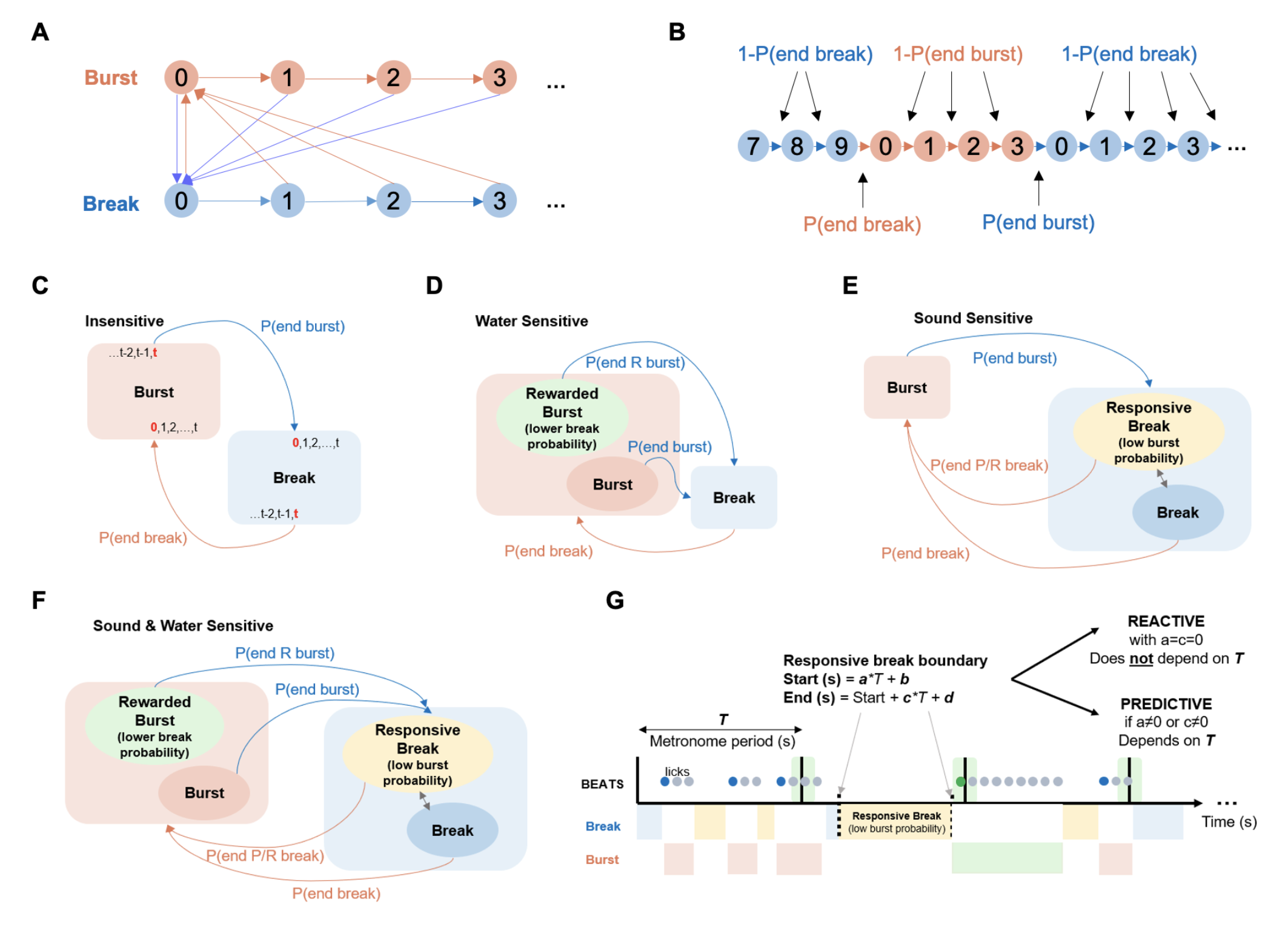
Modeling approach, with candidate strategies represented as Markov processes. **A)** A graph of the Markov process. The two branches correspond to lick bursts (red) and breaks (blue), each modeled as a succession of process states, one for each time bin of the data (10 ms bins used here). Transitions are either forward along the branch or back to the state corresponding to time 0 of the other branch. Thus, the termination of a lick burst is followed by a sequence of Break states, and the termination of a Break is followed by a sequence of Burst states. **B)** A schematic illustrating a trial decoded into the process states of the Markov process shown in A. A Break lasting 10 time steps is followed by a lick burst lasting 4 time steps, followed by another Break. All transition probabilities depend only on the current process state. The overall probability of this segment is the product of these transition probabilities. **C)** The Insensitive model assumes an alternation between the Burst and Break states with no dependence on external events. The transition probabilities are time-invariant. **D)** In the Water Sensitive model, the transition probabilities of the Burst state depend on whether the Burst is rewarded, as determined from the actual time course of the trial. **E)** In the Sound Sensitive model, transition probabilities of the Break state depend on the time since the last metronome beat. The tempo-dependence of the start and end of these “Responsive Breaks” determine whether the model is Reactive or Predictive. **F)** The Sound & Water Sensitive model combines the features of the Sound Sensitive (Responsive Breaks) and Water Sensitive (Burst states depend on reward) models. **G)** Schematic representation of the traversal through states in the time-dependent models of F. A hypothetical lick pattern is shown with licks in gray and the lick burst onset in blue if unrewarded, and in green if rewarded. Below the black lines representing the metronome are how the Sound & Water Sensitive models would parse the state that the animal is in, with the two types of Break states in blue (normal) and yellow (responsive), and the two types of Burst states in red (unrewarded) and green (rewarded). The boundaries of the Responsive Break are determined (as in Eq. 1 of the main text) by four parameters: *a, b, c,* and *d*. The responsive break is Reactive if *a* and *c* are both set to 0, or Predictive if *a* or *c* are nonzero.

The crucial assumption in all models is that the transition from one process state to the next one depends only on the current process state (although it may vary in time). This assumption makes the process into a Markov process. Given the Markovian assumption, the overall probability of an observed trial is simply given by the product of the probabilities of the transitions that occurred at each time step during the trial (Fig. 3A). The graph in Fig. 3A has one important consequence: successive durations of Bursts and Breaks should be independent of each other. Our data closely approximate this assumption (see Methods and Extended Data Figs. E1 and E2).

To conceptualize how the model works in practice, let us consider the intuitive case of human 1-to-1 tapping. The stimulus and tap sequence can each be discretized to produce a time series that represents the state of the external stimulus (beat or no beat) and the behavior (tap or no tap, corresponding to the Burst and Break states in Fig. 3A). Suppose that the time step of the model is 1 ms. A trial in which the metronome rate is 1 Hz would have a beat once every 1000 time steps. A human tapping to this metronome would also typically produce a tap once every ∼1000 time steps. A model that assumes a constant tap probability, irrespective of external events, would have a probability of 0.999 of progressing along the Break branch (and therefore its probability to generate a tap at any time point is 0.001). Since humans produce single taps, the probability to progress along the Burst state is 0 (and therefore the probability of exiting the initial Burst state would be 1). This model would produce single taps with inter-tap intervals that are widely distributed (having a geometric distribution with a mean of 1 s). However, such a model would be a poor fit to human tapping since human taps are concentrated around beats and occur with an inter-tap intervals of about 1 s.

By allowing the transition probability from no-tap → tap to be extremely low immediately following a beat, but then jump to a larger value at some point in the inter-beat period, it is possible to better capture human behavior. Importantly, the times at which these changes in probability occur reflect whether the process is predictive. For example, if this jump in tapping probability is constrained to occur at a fixed time interval following the previous beat, the model would be ‘reactive’ – reflecting the fact that tap times are not sensitive to the expected timing of the next beat. On the other hand, if the jump in transition probability occurs at a time point determined by a function of the beat interval, for example, jumping to a high no-tap → tap transition probability after 99% of the beat interval has elapsed, the model would generate tap patterns that are predictive, locking with a near-zero or negative mean asynchrony to the beat for all tempi. Therefore, even in the simple case of human 1-to-1 tapping, the beat rate-dependence of the parameters linking behavior to the external stimulus is the key diagnostic indicator of predictive synchronization.

Crucially, the modeling framework in Fig. 3 is not limited to the 1-to-1 beat to action relationship typical of human synchronization behavior. The temporal structure of the transition probabilities can still reflect predictive synchronization by being tempo-dependent, even with the 1-to-many lick bursts typical of rat behavior. We used this approach to construct models representing a range of candidate strategies that could have produced the observed data, with the specific aim of being able to quantitatively assess whether latent predictive synchronization is present in the lick patterns produced by rats performing this task.

The simplest possible strategy available to the rat is to ignore the stimulus entirely and produce lick bursts at random. We model this strategy by assuming constant transition probabilities between the Burst and Break states (a simple Markov chain with the state graph of Fig. 3A and time-invariant transition probabilities). Since the transition probabilities do not depend on external events, this is the **Insensitive** model (Fig. 3C).

The next model we tested is a **Water Sensitive** model (Fig. 3D), designed to accommodate the observation that rewarded lick bursts were longer in duration than unrewarded lick bursts. In the Markov framework, this can be expressed by postulating two sets of transition probabilities for a lick burst, one for **Rewarded Bursts** and another for unrewarded bursts. Like the Insensitive model, the Water Sensitive model consists of alternations between Break and Burst states, where the Burst state can be of either type depending on whether it is rewarded or not (as determined by the timing of the first lick of the burst). Note that this model does not assume any sensitivity to the metronome beat, and in fact is reduced to the insensitive model when the transition probabilities for the rewarded and unrewarded bursts are identical. The insensitive model is therefore ‘embedded’ within the water-sensitive model; this is a key feature in the construction of all models, making it possible to directly compare their likelihoods of producing the observed data.

The next class of models consists of two ***Sound Sensitive*** models. These models incorporate a sensitivity to the metronome that may be either **Reactive** or **Predictive** depending on the tempo-dependence of the parameters. The main experimental observation (Fig. 2G) is the elevation of the lick rate close to metronome beats and its reduction between them. We conceptualized this as a strategy in which the rat lowers its lick burst rate during a time interval that starts shortly after a metronome beat and ends before the next one (Fig. 3E). We consider this period of lower burst rate as the response of the rat to the metronome beat, and therefore call the breaks during that time ‘responsive breaks,’ as opposed to the normal breaks that occur outside of this window. During responsive breaks, we expected the transition probabilities from Break to Burst to be smaller than during normal breaks, presumably reflecting a deliberate suppression of licking. This responsive break can be independent of tempo (with fixed start and end times) reflecting a reactive strategy, or tempo-dependent, indicating predictive behavior.

The last class of models, the ***Water & Sound Sensitive*** model (Fig. 3F) combines the Water Sensitive and Sound Sensitive models. Transition probabilities for lick bursts depend on water rewards and transition probabilities for breaks depend on the metronome beats. The tempo dependence of the start and end of the responsive breaks determine whether the models are **Water Reactive** or **Water Predictive**.

In all sound sensitive models, the times (specified relative to the most recent metronome beat) at which transition probabilities shift from normal to responsive and back are denoted by t_start_ and t_end_. The dependence of these times on the period of the metronome is our main argument for the presence of latent predictive synchronization in rats. We parametrized the responsive window by the following equations:

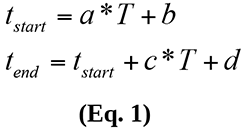

where *T* is the metronome beat period (in seconds), the parameters *a* and *c* are unitless fractions of the beat interval (bounded between 0 and 1), and the parameters *b* and *d* are in seconds (constrained to be between −0.5 and 0.5). To prevent the start and end of the *Responsive Break* from exceeding the entire beat interval, *t_start_* and *t_end_* were constrained by setting values that were less than zero or larger than *T* to 0 and *T*, respectively, and setting t_end_ values less than *t_start_* to *t_start_* (resulting in no responsive break in these cases).

The distinction between Reactive and Predictive models is based on these equations. In Reactive models, the parameters *a* and *c* were constrained to be zero, so that the Responsive Break begins *b* seconds after each beat and ends *b+d* seconds after the beat, regardless of tempo. Note that such responsive breaks do not require prediction of future beats. We consider such a model to reflect a reactive strategy. When *a* and *c* are nonzero, the responsive window becomes dependent on the beat period. Nonzero values of *a* and *c* therefore imply the existence of an internal prediction of when the next beat will occur, reflecting a predictive strategy.

The different strategies in Fig. 3 can be quantitatively compared to determine which one best explains the observed behavioral patterns. Briefly, the six models were fit to the data by discretizing trials into 10 ms bins and assigning a process state to each time bin based on whether the animal was in a lick Burst or Break (all models), whether that Burst was rewarded or not (Water Sensitive, Water Reactive, & Water Predictive models only), and whether the Break was a Responsive Break or not as deemed by the *a, b, c,* and *d* parameters of the model being optimized (Reactive, Predictive, Water Reactive, & Water Predictive models only). We parameterized the Burst and Break transitions using a small number of parameters (see Methods for details). The log probabilities of all the transitions observed in the data given each model were summed over all time bins and trials to produce an overall log-likelihood of that model. These log-likelihoods were maximized over the parameters for each model. The optimized log-likelihoods of embedded models were then compared using χ^2^ tests, allowing us to test the null hypothesis that a=c=0.

### Rats predictively time their licking

Table 1 and Fig. 4A compare the log-likelihoods of the different models, using the Insensitive model as a reference. Water sensitivity substantially improved model fits compared to the corresponding models that did not include water sensitivity (first three rows of Table 1), reflecting the rats’ tendency to produce longer lick bursts during water reward. Sound sensitivity also substantially improved the fit to the data (rows 4-5 of Table 1). Most importantly, incorporating a dependence of the responsive window on tempo further substantially improved the quality of the data fit (rows 6-7 of Table 1). The Water Predictive model was by far the best fitting model, outperforming the second best model, the Water Reactive model, by a highly significant 115.6 log-likelihood units (χ^2^(2)=231, p<10^-50^).

**Figure 4.**
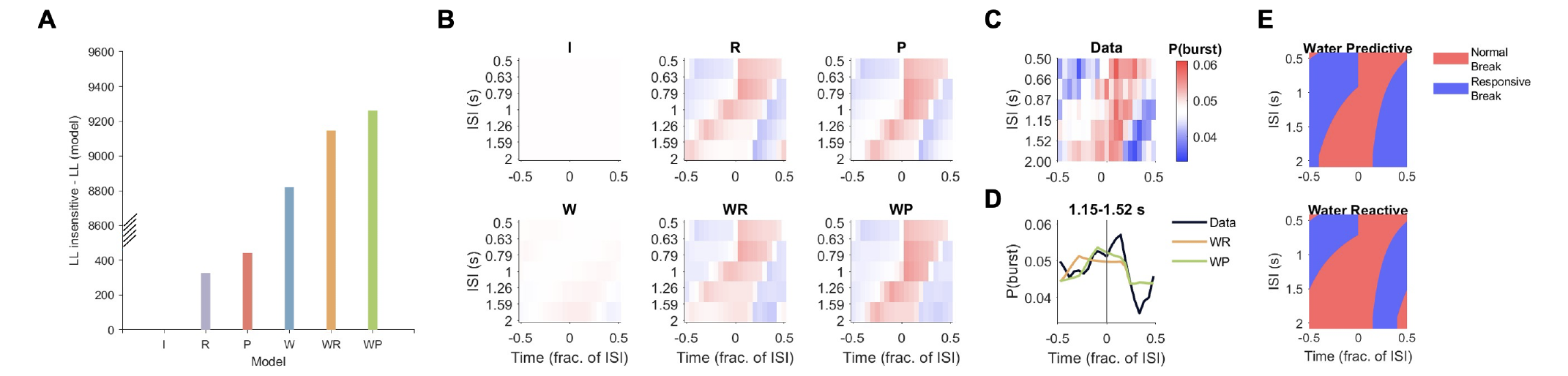
Lick patterns are best explained by a predictive model. **A)** Log-likelihoods of the candidate models, referenced to the Insensitive model. Note the break in the y-axis before the water sensitive models. **B)** Each panel depicts probability of lick bursts split by tempo bin based on the six models tested (I: insensitive, R: reactive, P: predictive, W: water sensitive, WR: water+reactive, WP: water+predictive). **C)** Observed licking rates in five tempo bins (same as Fig. 2D, reproduced here to ease comparisons). **D)** Data versus the WR (orange) and WP (green) models for an intermediate tempo bin. Note that the WP model better captures the timing of the rise and fall in lick burst probability compared to the WR model. **E)** The temporal extent of the Responsive break period (blue) across tempi, based on the optimized *a, b, c,* and *d* parameters. The normal break, characterized by higher probability of emitting a lick burst, is shown in red. The WP model (top) allows for shortening of the normal breaks relative to the WR model (bottom), improving the fit to the data in panel C.

**Table 1.** Summary of model comparisons.

| Comparison | $\Delta$ LL | $\Delta$ df | p-value |
| --- | --- | --- | --- |
| I vs. W | 8819 | 1 | 0 |
| R vs. WR | 8819 | 1 | 0 |
| P vs. WP | 8819 | 1 | 0 |
| I vs. R | 327 | 3 | $10^{-140}$ |
| W vs. WR | 326 | 3 | $10^{-140}$ |
| R vs. P | 116 | 2 | $10^{-50}$ |
| WR vs. WP | 116 | 2 | $10^{-50}$ |

We used the optimized parameter values to compute the Burst probabilities around beats for each model tested (Fig. 4B) for comparison with the observed data (Fig. 4C). As expected, the Insensitive model shows no structure at all in the timing of lick bursts, and serves here as the baseline reference for the more complex models in which it is embedded. The estimated burst rate of the Insensitive model, and indeed all models, is identical to the average burst rate of the data (µ=0.048 per 0.1*ISI bin, Fig. 4B-C). All other models, including the Water Sensitive model that has no direct dependence on the stimulus, showed modulation of the rates of lick bursts around beat times. However, the introduction of sound sensitivity through responsive breaks substantially improved the fit to the data.

To illustrate the better fit of the predictive model to the data, Fig. 4D shows the probability of lick bursts around metronome beats computed from the (best fitting) Water Predictive model compared to the (second best fitting) Water Reactive model for metronome periods between 1.15 and 1.52 s. The predictive model captures very well the time at which the lick burst rate increased before the beat (green line), while the reactive model (orange line) shows an increase in the lick burst rate that is clearly too early compared with the data. Thus, the reactive model, with its fixed duration window for responsive breaks, is unable to account for the period-dependence of the observed changes in lick burst rate.

The optimized parameters provide further information on the strategies used by the rats to solve the task (Table E1 in Extended Data; see Methods for a detailed description of parameters). The rewarded Bursts were indeed longer (smaller exit probability) than unrewarded Bursts. Also as expected, the Responsive Breaks tended to be longer than normal Breaks. Fig. 4E visually compares the optimized Responsive Break periods for the Water Predictive and Water Reactive models (determined by the *a, b, c,* and *d* parameters in Eq. 1). The beginning of the Responsive Break period (the transition from red to blue at positive phases) is identical in the two models and starts 300 ms after each beat, strongly suggesting that the rats indeed started their suppression of licking in reaction to the beat. However, the end of the Responsive Break (the transition from blue to red) differs in the two models. While the blue area has a constant duration in terms of time (and therefore a longer extent in terms of phase for fast vs. slow tempi) for the Reactive model, its temporal extent is longer at slower tempi in the Predictive model, indicating longer suppression of licks (blue) at slower than at faster metronome rates. This longer suppression period reflects an internal prediction of when the next beat is going to occur.

### Rats shorten their lick bursts at faster tempi

The finding that responsive breaks were tempo-dependent raises two related questions. First, are there any other model parameters that are tempo-dependent? And second, is the dependence of the responsive break window on tempo indeed linear, or are there non-linearities in this dependence?

To address these questions, we ordered trials by tempo and split them into five groups, with approximately equal numbers of trials in each tempo group. We then fitted all six models onto the trials in each tempo group. For the Sound Sensitive models, we added the additional constraint that the *a, b, c,* and *d* values (Reactive, Predictive, Water Reactive, Water Predictive models) should result in t_start_ and t_end_ values that were continuous at the boundaries of neighboring tempo groups, based on the intuition that the animal should not change its strategy abruptly from one tempo to the next. We call this the 5-group model, while the model described in the previous section is the 1-group model.

Fig. 5 shows the optimized parameter values derived from this 5-group analysis. Fitting models to data split by tempo improved the model fits substantially for all models (**I:** χ^2^(12)=619, p<10^-50^; **W:** χ^2^(16)=446, p<10^-50^; **R:** χ^2^(16)=701, p<10^-50^; **P:** χ^2^(24)=628, p<10^-50^; **WR:** χ^2^(20)=527, p<10^-50^; **WP:** χ^2^(28)=455, p<10^-50^), indicating that indeed other aspects of rat behavior were modulated by tempo. We compare first the start and end times of the responsive breaks from the 5-group model with those from the 1-group model (Fig. 5A-B). In spite of the larger freedom available to the 5-group model in setting these boundaries, the two models determine surprisingly similar responsive break periods as a function of tempo, suggesting that the linear assumption of the 1-group model is actually rather closely followed by the data.

**Figure 5.**
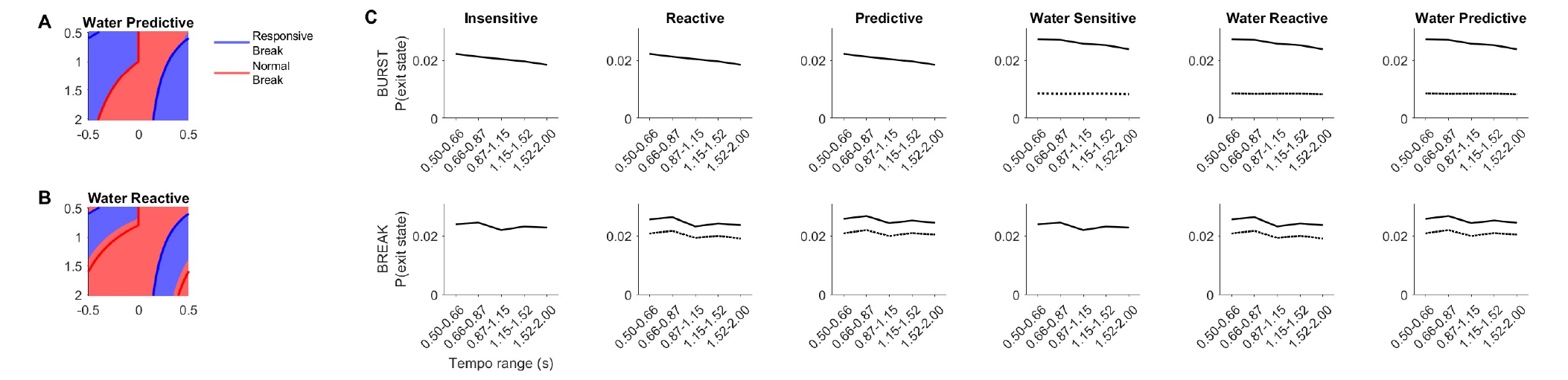
Optimized parameter values reveal further sensitivity to tempo. **A)** Optimized temporal boundaries of Responsive Break periods for the 5-group Water Predictive model. The lines depict the temporal boundaries of the 1-group model for comparison. **B)** Same as A, but for the Water Reactive model. **C)** Optimized probabilities of exiting the Burst (top row) or Break (bottom row) state for each of the six models (columns).

The high similarity between the temporal boundaries of the 5-group and 1-group Water Predictive models indicates that the improvement in fit was primarily due to the rats adjusting their transition probabilities by tempo. Figure 5C shows the probability of exiting the Burst state (top row) and Break state (bottom row) for the different models in the five tempo bins. Most of the transition probabilities did not depend strongly on tempo. These included the transition probabilities out of a responsive break (Fig. 5C bottom, dashed line), normal break (Fig. 5C bottom, solid line), and rewarded burst (Fig. 5C top, dashed line). On the other hand, the probability of exiting unrewarded Burst state (Fig. 5C top, solid line) showed a clear tempo-dependence, decreasing almost linearly with longer beat periods. This tempo-dependence was present in all models. We interpret this finding as reflecting a behavioral strategy where the rats shortened their unrewarded lick bursts at faster tempi in order to produce the minimal break duration required to get reward. Such changes in behavior may improve success rate, but are not necessarily predictive. Thus, these results indicate that the rats adjusted their behavior to metronome rate using a mixture of predictive and non-predictive strategies.

## Discussion

This study presents a nuanced exploration of behaviors that may have previously been dismissed as useless for studying auditory-motor synchronization. We make here two conceptual advances: first, the introduction of the notion of latent predictive synchronization, together with the most concrete evidence to date that rats are capable of such predictive synchronization in a range of tempi compatible with human behavior. Second, we introduce a flexible modeling approach that enables rigorous comparison of candidate strategies accounting for an observed behavior during a task. In the context of rhythmic synchronization to metronomes, this approach resolved the hurdle posed by the fact that the rats showed a 1-to-many beat-to-action synchronization behavior, and allowed us to explicitly demonstrate that rats adopted a predictive strategy in synchronizing their licking to metronomes.

Our study shares similarities with previous studies of auditory-motor synchronization in rats, but also complements them in crucial ways. One previous metronome synchronization study in rats [7] employed lever-pressing and revealed that rats produce a behavior that was clearly tempo-matched to the stimulus. However, that study did not provide evidence that the synchronization was predictive. Here we go further by demonstrating that rats are producing a behavior that is unequivocally predictive. Specifically, we show that rats terminated a period of suppressed licking between beats as a function of the metronome period, showing an anticipation of the next beat. It is worth noting that the reward window differed greatly between the two tasks. Katsu and colleagues employed a reward window that included a large fraction (>70%) of the inter-stimulus interval, with most of the reward window occurring *after* the beat. With such a reward window, rats would not gain much by adopting a predictive strategy. In our task, the window for reward was short (+/- 7.5% of the inter-stimulus interval) and symmetric around beat onsets, so that anticipation was presumably advantageous for obtaining reward. These differences in task design may also be responsible for the positive mean asynchronies observed by Katsu and colleagues, in contrast to the negative mean asynchronies we observed here (see Fig. 2F).

A study that explored spontaneous synchronization to beat in rats listening passively to music [12] found head movements that were locked to beats. However, that study did not demonstrate that these head movements included a predictive component. Curiously, these synchronized head movements were only present during the first experimental session, which is suggestive of startle-like reactions that habituate rapidly. Similar spontaneous and seemingly involuntary facial movements locked to a rhythmic auditory sequence have recently been reported in head-fixed mice as well and were extinguished when associated with an aversive stimulus (see Fig. 1C in [26]). To the extent that these involuntary responses share neural circuitry with the networks responsible for voluntary auditory-motor synchronization, startle responses may still prove to be an important component of the synchronization process. Our modeling approach may in fact help quantify the subtle ways in which a spontaneous behavior in the absence of a task may still be driven by an external stimulus. For example, our framework could be applied to test whether such involuntary movements are executed predictively when the auditory rhythm is predictable – reflecting a spontaneous form of predictive synchronization – or whether they follow the salient stimulus events with a short but fixed latency. Such models would therefore link task-free approaches [12] with studies of voluntary auditory-motor synchronization such as this one.

To our knowledge, our implementation of a Markov process that is responsive to external events provides a novel framework for modeling sensorimotor synchronization. It differs from existing models of sensorimotor synchronization, which have typically taken dynamical systems [27, 28], Bayesian [29], or a combination of the two approaches [30]. Our approach conceptually takes a step backward, engaging directly with the observed behavior rather than with mechanisms (as in dynamical models) or with normative accounts (as in Bayesian models). Importantly, our models and their parameters are directly interpretable, allowing for the identification of elements of both predictive and non-predictive strategies involved in producing a synchronization behavior. We believe this approach can lead to a more nuanced insight into human sensorimotor synchronization as well. Human synchronization is often taken for granted despite high variability between individuals [22, 23] and the existence of high and low synchronizers [31, 32]. We conjecture that humans too likely employ a mix of predictive and non-predictive strategies while synchronizing, with this mix depending on the individual, the stimulus, and the effector producing the action. Our modeling approach breaks conceptual ground by shifting from a view of the 1-1 beat-to-tap correspondence as reference for all aspects of sensorimotor synchronization, which is an idealization even when considering only humans, to one that is based on unbiased modeling of the observed behavior. In consequence, this approach enables the extraction of general strategies that may be employed by any species, human or nonhuman, under any stimulus context. Given its generality, this approach could perhaps also be used to model a neuron’s probability of spiking throughout a beat interval to differentiate responses evoked by, versus those entrained to, rhythmic sensory input [33].

Finally, our findings have implications for theories of the evolutionary origins of rhythm perception and auditory-motor synchronization. They provide strong evidence against the vocal learning hypothesis [13] by demonstrating that the rat, a non vocal learner, can be trained to synchronize predictively to a metronome. Alternatively, the gradual audiomotor evolution hypothesis suggests that the capacity for rhythmic entrainment developed gradually across primates, peaking in humans but present to a limited degree in non-human primates [14]. Our results support this hypothesis, and further extend it to cover a wider range of mammalian species. In consequence, our findings build a potential bridge between the extensive electrophysiological work done in rodent models on timekeeping during relatively simple tasks [34-38], with the growing insight coming from in-vivo recordings of nonhuman primates synchronizing to metronomes [15, 16, 39, 40]. Beyond their biological and evolutionary implications, this work demonstrates the feasibility of the study of beat perception and synchronization in rodent models, and facilitates direct comparisons of the dynamics of sensorimotor entrainment as they vary across species in the animal kingdom.

## Methods

### Behavior

#### Animals

All procedures were performed according to protocols approved by the Animal Research Ethics Subcommittee of the City University of Hong Kong. Eight female rats (Wistar, 250-280g) with normal acoustic startle responses were tested. Rats were on a schedule of 5 days of testing (2 sessions/day at least 4 hours apart), during which water was a positive reinforcer, followed by 2 days of ad lib water. Water bottles were removed 16 hours before the first testing session of the week. During testing, reward amounts were adjusted such that rats received no less than 8 ml per day. If a rat’s performance was especially poor, the required amount of water was delivered in the rat’s home cage.

#### Behavioral Setup

The behavioral setup consisted of a custom-built cage with one water spout controlled by a solenoid valve, a speaker, and a white LED placed overhead. A simple transistor circuit was constructed that would pass a small voltage whenever the rat made contact with the spout, and a TDT RZ6 real time processor (Tucker Davis Technologies, Florida) was used to detect licks and open the water spout if required with < 1 ms delay. The TDT processor was controlled using custom code written in RPvdsEx, and experiments were run using custom-written Python code in the Spyder environment from a Windows PC. The behavioral box was placed inside an acoustically shielded chamber, and rats were placed into the box one at a time and monitored by the experimenter using a web camera.

#### Training

Training was conducted in three stages. In all stages of training and testing, rats performed two ∼16 minute sessions per day. In stage 1, any licks to the water spout were rewarded with a small drop of water, and through this the animals were familiarized with licking the spout for reward. This stage also enabled us to measure the natural lick rate of the rats (roughly 7 Hz; see Extended Data Fig. E4A). Despite several attempts, we were not successful in training the animals to slow down the rate of their licks.

The second stage of training was a “T-on, T-off” task, which was designed to train rats to 1) associate sound (white noise) with reward availability during the sound on period (T), and 2) break contact with the spout during silence for a minimum amount of time (T) before reward would become available again. In this task, all licks during “T-on” would be rewarded, and any licks during “T-off” would be unrewarded and additionally would reset the “T-off” period. Thus, during “T-off,” for sound (and thereby, reward) to become available again, the rat had to avoid touching the spout for at least T seconds. The rats quickly understood this and adjusted their inter-lick intervals (ILIs) to match T (Extended Data Fig. E4B). Thus, the rats were capable of suppressing their licks, often by biting the spout or moving the head away from the spout, for a flexible amount of time that was clearly stimulus-specific, even if they could not slow down their natural lick rate. A single value for T was selected and presented continuously to the animal during a given session.

In the third and final stage of training, rats were introduced to the metronome task with a more generous reward window that was gradually tightened (± 20%, 15%, 10% of ISI). Altogether, training lasted roughly 3 weeks, with roughly half of this time spent in the final, third, stage.

#### Variable ISI Metronome Task

Each ∼16 minute session consisted of approximately 45 trials, where each trial consisted of 16 isochronous white noise bursts. The inter-stimulus interval (ISI) for each trial was drawn at random from a log-spaced uniform distribution between 0.5 s and 2 s such that trials were equally probable in the two octaves of possible ISIs spanned by the experiment. The duration of the white noise burst was 7.5% of the ISI and represented the latter half of the time window during which reward was possible. To trigger the spout to open and deliver a small drop of water reward, a lick needed to be within ±7.5% of the ISI from a beat onset *and* needed to be separated from the preceding lick by at least 450 ms. This ensured that licking continuously would not be rewarded. Each trial in a session lasted between 8 s and 32 s (16 x ISI) and was separated from the next trial by 4-5 s (drawn randomly from a uniform distribution) of silence. A 1000 ms 5 kHz tone was played as soon as the 16th beat ended to signify the end of the trial. As a further incentive, the size of the water reward was incrementally increased for each successful burst in a trial and reset to the baseline amount at the start of the next trial. On occasion, rats were unmotivated during a session and would not engage with the spout. Thus, between 18 and 22 sessions were collected per animal. During each ∼16 minute session, rats typically performed ∼45-50 trials.

#### Data Analysis

Trials containing no licks were discarded from analysis. As described in *Training*, we could not train rats to slow down their natural ∼7 Hz lick rate, but we were able to train rats to suppress licking for specific time intervals driven by an auditory stimulus (Fig. 3B). Thus, only the first lick of each lick burst, which needed to be separated from the preceding lick by at least 450 ms, were analyzed because they were considered a reflection of the animal’s intention to synchronize with the metronome to receive reward. Burst asynchronies were calculated by subtracting from each burst time the nearest beat onset and dividing the result by the ISI of that trial. Burst asynchronies thus ranged between −0.5 and +0.5, where 0 asynchrony would be perfectly synchronous with the beat onset.

As a control to test whether rats had found and adopted some strategy that did not require them to listen and adjust their behavior based on the auditory metronome, a conservative simulation was performed. The simulation left the temporal structure of the rats’ licks intact and simply assumed that the behavior was in response to a different metronome tempo. Simulation ISIs were drawn at random from the same distribution as the true ISI experienced by the animal in a trial. If a simulated ISI was larger than the true ISI for a trial, then fewer than 16 beats could be analyzed based on the simulation, and in these cases, the real data were truncated so that the same number of beats were being analyzed between real and simulated data. This simulation was run 100 times, and the data shown in Fig. 2G compare averaged simulation data with real data truncated where necessary to match the duration of simulation trials.

### Markovian Model

A conceptual description of the Markovian modelling approach is provided in section 2 of the Results and in Fig. 3. The technical details are provided below.

#### Model structure

Rat behavior was analyzed at a resolution of 10 ms. The graph of the model (Fig. 3A) illustrates the alternation through time between two behavioral states, described by the two “branches” of the graph: Burst and Break. Within each branch, each state has two possible transitions: either forward to the next state within the branch, representing an elongation of the current behavioral state, or to the first state of the other branch, representing the end of a lick burst (and the start of a break), or the end of a break (and the initiation of a lick burst). The branch lengths should be considered as infinite, but in practice rats reached only a finite set of these states, since lick bursts were typically shorter than 1s (100 states) and breaks could not be longer than 32 s (3200 states, trial duration of the longest trial). For simplicity, we set a maximum duration that the rat could spend in any behavioral state of 10 s (1000 states). The transition probabilities of the model depend on the current state only, and not on past history, but were allowed to potentially vary based on external events in temporal proximity to the current state (e.g. metronome beats and water reward), depending on the model.

The time-dependence of the transition probabilities along the Burst branch of the process is induced by water reward. If a lick burst is initiated within the rewarded window, the lick burst is rewarded. Note that whether a burst is rewarded or not cannot be modeled by a probabilistic transition, since the decision of whether or not to reward a burst is deterministic based on the timing of the lick burst. In the water-dependent models, we assume separate sets of transition probabilities for unrewarded and for rewarded bursts.

The time-dependence of the transition probabilities along the Break branch of the process is induced by metronome beats. In the sound-sensitive models, we posited a ‘responsive window’ during which lick burst probability decreased. We modeled this by assuming two separate sets of transition probabilities along the Break branch, one used during normal Breaks and the other for the Responsive Breaks. The set of parameters used was determined by the time of the specific transition. Thus, during a single Break, transition probabilities could shift from normal to responsive and back. In Fig. 3G, this occurs for the marked break in the middle of the scheme – it starts before the beginning of the responsive window but extends into it.

The Insensitive model, where all transition probabilities are time-invariant, can generate any distribution of Burst durations and Break durations by setting the transition probabilities appropriately. For example, if L is the random variable of lick burst durations, the probability of going from state n of the Burst branch to the initial Break state is *P*(*L*= *n*| *L n*) (the probability of the lick burst being of length n exactly, given that it is at least n). It can be shown that the distribution of the duration of lick bursts generated by a model with these transition probabilities will simply be that of L. It follows that the model is highly expressive in that it can fit many different distributions.

There is nevertheless one important characteristic of the data generated by the model – in the time-invariant case, the durations of successive Bursts and Breaks are statistically independent. In other words, knowing the duration of a lick burst does not change the distribution of the next break, or of the Burst after that Break, and so on. This independence property is not necessarily kept in the time-varying case, since external events may induce correlations between the durations of successive Bursts and Breaks. One potential source of small correlations in the data are slight differences in behavior between rats. For example, if one rat tends to overall produce shorter episodes of Bursts and Breaks, and another rat tends to produce longer ones, the overall data may show weak correlations. In our data, the fraction of explained variance in the durations of successive lick bursts and breaks was r^2^=0.0037, and is significant because of the large number of Burst and Break pairs in the data (N=74,279; see Extended Data Fig. E1). However, since the correlation is so small, and because we believe they were induced by the small inhomogeneities in the data described above (see Extended Data Figs. E2), we proceeded with the Markovian assumption.

#### Model parameters

Separate optimization of every transition probability from any state of the process to its two targets would yield a few thousand parameters to estimate, but would result in low statistical power to discriminate between models. Therefore, we limited the testing of the models to only a few families of parametric distributions.

For Bursts, we tested the geometric distribution *`P(L=n) = p(1-p)`^(n-1)^* as well as *`P(L=n) = np`^2^`(1-p)`^(n-1)^*, which yields a probability distribution that peaks at one side or another of *`1/p`*, and decreases asymptotically like the geometric distribution for large n. Both distributions were parametrized by *`p`*, the ‘exit probability’ (or asymptotic exit probability in the second case) from the branch and into the initial state of the break branch. In both cases, it is possible to compute analytically the transition probabilities along the Burst branch of the model: they are *`1-p`* (independent of position along the branch) and *`(1-p)`^2^ `/ [1 – (1-1/n)p]`* for the respective distributions, with the exit probabilities being their respective complements.

For lick burst durations, the geometric distribution was invariably a better fit to the model. In the water-sensitive models, we specified two separate parameters for the lick burst durations, one for the unrewarded and one for the rewarded licks. Thus, lick bursts durations were modeled as having geometric distributions, with rewarded licks turning out to have a smaller exit probability at each position (longer lick bursts) than unrewarded lick bursts (e.g. Fig. 5).

To model break durations, we had to implement the condition that breaks had a minimal duration of 450 ms (45 steps of the process). Due to a small number of <450 ms breaks in the data (presumably at the beginning and end of trials), we parametrized the first 45 steps of the break branch by a separate, fixed exit probability. After optimization, this exit probability turned out to be indeed small, accounting well for the deadtime of the distribution of breaks. Beyond these initial 45 steps, transitions in the break branch could be normal or responsive, depending on the time of the state relative to the last metronome beat. Normal and responsive break transitions were parametrized by two separate exit probabilities.

Thus, probabilities along the break branch of the model could switch seamlessly from one value to the other, in case a break that started during the normal window extended into the responsive window, or vice versa. This parametrization resulted in a complex distribution for break durations, with a 450 ms deadtime of durations that had extremely low probability of occurring, followed by an approximately geometric distribution of durations. The start and end times of the responsive windows were parametrized by linear functions of the metronome period (Eq. 1 in the main text).

These parametrizations resulted in models that had up to 5 transition probability parameters (exit probabilities for unrewarded and for rewarded Bursts; exit probability for the partial deadtime of Breaks; exit probabilities for the normal and responsive Breaks). In addition, the sound-sensitive models had either 2 (in the Reactive case) or 4 (in the Predictive case) additional parameters describing the linear dependence of the responsive window on the metronome period. Table 2 summarizes schematically the parameter structure of each of the six models tested. Table 3 in Extended Data additionally fills in Table 2 with the optimized parameters from the main analysis, and Fig. 5 in the main text shows how these parameters varied with tempo based on the 5-bin, tempo-resolved model.

**Table 2.** The parameter logic of all models. The first five columns specify the possible transition probability parameters. These are always parametrized by the exit probabilities, pointing from a state of one branch of the chain to the first state of the other branch (see Fig.3A). NBu and RBu are are exit probabilities from the states along the Burst branch. In the water-sensitive models, there are separate transition probabilities for unrewarded (NBu) and rewarded (RBu) Bursts. In the Insensitive and the sound-sensitive models, these two probabilities are equal. DBr is the exit probability during the first 45 time steps (450 ms) of a Break. In the sound-sensitive models, there are separate transition probabilities for normal (NBr) and responsive (RBr) Breaks. In the Insensitive and Water models, there is no responsive window, and thus the two probabilities can be considered as being set to be equal. Columns a-d are the parameters of the beginning and end of the responsive window (as defined in Eq. 1 of the main text). They can be considered as being equal to 0 in the Insensitive and Water models; the slope parameters, a and c, are set to 0 in the Reactive models.

| MODEL | NBu | RBu | DBr | NBr | RBr | a | b | c | d |
| --- | --- | --- | --- | --- | --- | --- | --- | --- | --- |
| Insensitive | X | = | X | X | = | 0 | 0 | 0 | 0 |
| Water | X | X | X | X | = | 0 | 0 | 0 | 0 |
| Reactive | X | = | X | X | X | 0 | X | 0 | X |
| Predictive | X | = | X | X | X | X | X | X | X |
| Water Reactive | X | X | X | X | X | 0 | X | 0 | X |
| Water Predictive | X | X | X | X | X | X | X | X | X |

#### Fitting the models

To estimate these parameters, all trials were first decoded, with each time bin of each trial assigned as either part of a Burst or Break (Fig. 3B). Given the parameters of the model being fitted, the specific type of behavioral state was determined (unrewarded vs. rewarded Burst, and normal vs. responsive Break). Based on this decoded representation of a trial given a specific model, the transition probabilities for all the transitions that actually occurred in the data were computed. We then summed the logarithms of these probabilities to get the overall log-likelihood of a trial given the model. The log-likelihoods were summed over all trials and all rats to give the overall log-likelihood of the data given a model with this set of parameters. The maximization of this overall log-likelihood was performed in two steps. First, for a given set of responsive window parameters, the transition probability parameters were optimized using the Matlab routine fmincon (matlab 2020a-2023a). The step was sufficient for the Insensitive and for the Water models, which did not have a responsive window. For the sound-sensitive models, the log-likelihood map as a function of the four (or two) responsive window parameters tended to have an uneven landscape of values, with multiple small local maxima. Therefore, for the sound-sensitive models, the transition probabilities were optimized for a grid of possible values of a, b, c, and d parameters, with b and d ranging independently from −0.5 to 0.5 s in 0.1 s increments, and a and c parameters ranging independently from 0 to 1 (fractions of the ISI) in increments of 0.1 for the Predictive models, and set to 0 for the Reactive models. The parameter values that had the maximum likelihood over this grid were selected for the optimal model.

To compare the goodness of fit of the six different models, we used log-likelihood tests for nested models. A nesting occurs when the parameters of one model can be reduced to those of another by setting some of them to predetermined values (see Table 2 for the nesting relationships between the models considered here). Importantly, the log-likelihood of nested models can be compared: the log-likelihood of the richer model will always be larger than that of the reduced model. Under the null hypothesis that the data are independent of the additional parameters, 2·ΔLL∼χ^2^(df), where df is the number of additional free parameters of the richer model [41]. We only compare nested models here.

## Extended Data

**Fig. E1.**
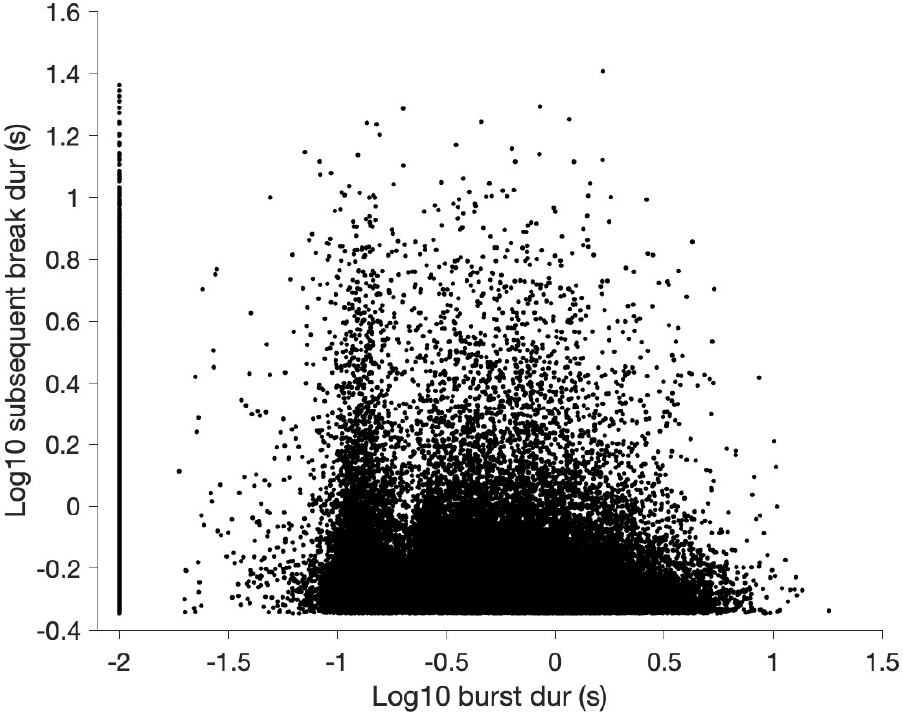
Joint distribution of burst durations and subsequent break durations. Due to both distributions having long right tails, their logarithms (base 10) are plotted in order to make the data more uniformly distributed. There is a significant correlation due to the large number of data points (r^2^ = 0.0037, p<10^-61^, N=74,279 pairs of bursts and breaks), but the coefficient of determination is extremely small. Single-lick bursts were assigned a nominal duration of 0.01 s.

**Fig. E2.**
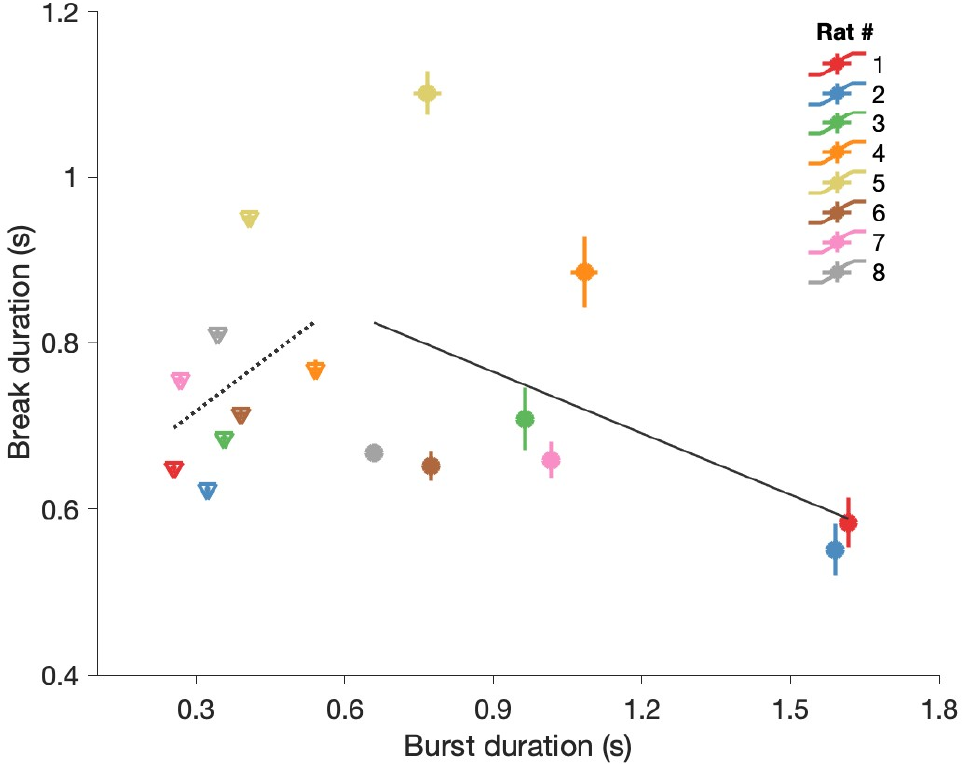
Average Burst durations and their subsequent Break durations for individual rats. We computed the averages separately for rewarded (circles) and non-rewarded (triangles) lick bursts. The gray lines are regression lines for the rewarded (solid line) and unrewarded (dotted line) licks. Error bars are SEMs. Note the clustering of points based on whether a lick burst was rewarded or not, and the differences between individual rats, particularly in their typical burst durations. The regular tendencies of these data may contribute to the small correlation we see between burst and subsequent break durations (Fig. E1).

**Table E1.** **Optimized parameter values for all models based on main analysis.** The five columns to the right of the MODEL column list the transition probability of exiting the given behavioral state (**NBu:** Normal Burst, **RBu:** Rewarded Burst, **DBr:** Deadtime Break, **NBr:** Normal Break, **RBr:** Responsive break). Final four columns are the a, b, c, and d parameters determining the temporal boundaries of the Responsive Break state. Probabilities are always specified for time bins whose duration is 10 ms.

| MODEL | NBu | RBu | DBr | NBr | RBr | a | b | c | d |
| --- | --- | --- | --- | --- | --- | --- | --- | --- | --- |
| Insensitive | 0.0200 | = | 0.0017 | 0.0232 | = | 0 | 0 | 0 | 0 |
| Water | 0.0254 | 0.0084 | 0.0017 | 0.0232 | = | 0 | 0 | 0 | 0 |
| Reactive | 0.0200 | = | 0.0017 | 0.0244 | 0.0205 | 0 | 0.3 | 0 | 0.5 |
| Predictive | 0.0200 | = | 0.0017 | 0.0251 | 0.0208 | 0 | 0.3 | 0.2 | 0.5 |
| Water Reactive | 0.0254 | 0.0084 | 0.0017 | 0.0244 | 0.0205 | 0 | 0.3 | 0 | 0.5 |
| Water Predictive | 0.0254 | 0.0084 | 0.0017 | 0.0251 | 0.0208 | 0 | 0.3 | 0.2 | 0.5 |

**Fig. E4.**
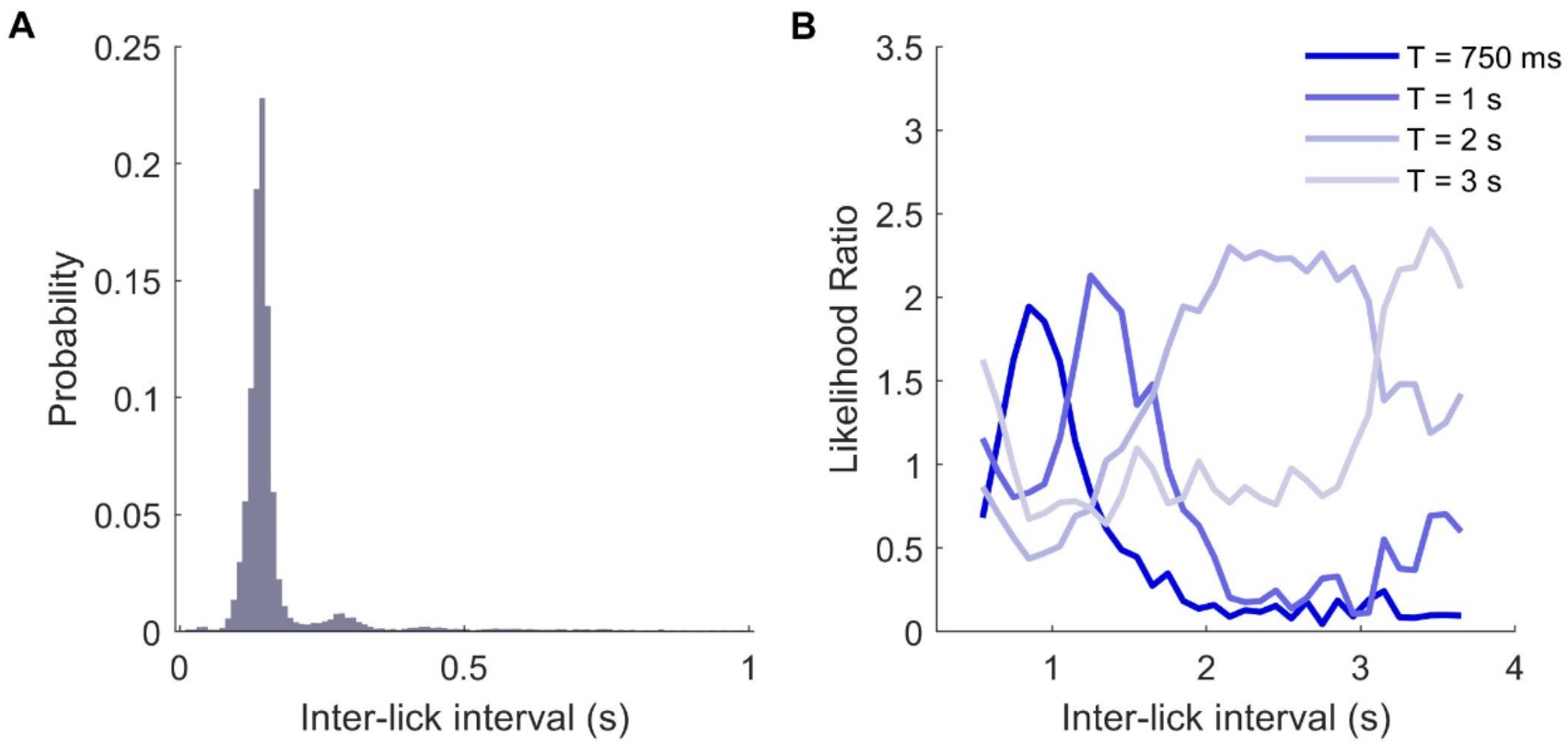
Rats show highly stereotyped licking patterns that can be flexibly switched on and off. **(A)** Inter-lick intervals (ILIs) while rats lick freely, pooled across animals and sessions. The peak is at ∼140 ms and corresponds to roughly 7 Hz. **(C)** ILIs in the T-on T-off task. Water reward was available only during T-on, and any licks during the T-off period were unrewarded and would furthermore reset the T-off period. Rats quickly adjusted their ILIs to match T. Y-axis is the likelihood of observing a given ILI for a condition relative to the probability of observing that ILI across all conditions pooled.

